# Fentanyl blockade of K^+^ channels contribute to Wooden Chest Syndrome

**DOI:** 10.1101/2025.01.17.633656

**Authors:** Aguan D. Wei, Nicholas J. Burgraff, Luiz M. Oliveira, Thiago S. Moreira, Jan-Marino Ramirez

**Author notes:** These authors contributed equally to this work (Aguan D. Wei, Nicholas J. Burgraff).

## Abstract

Fentanyl is a potent synthetic opioid widely used perioperatively and illicitly as a drug of abuse ^1,2^. It is well established that fentanyl acts as a μ-opioid receptor agonist, signaling through Gα_i/o_ intracellular pathways to inhibit electrical excitability, resulting in analgesia and respiratory depression ^3,4^. However, fentanyl uniquely also triggers muscle rigidity, including respiratory muscles, hindering the ability to execute central respiratory commands or to receive external resuscitation. This potentially lethal condition is termed Wooden Chest Syndrome (WCS), the mechanisms of which are poorly understood ^5–7^. Here we show that fentanyl directly blocks a subset of EAG-class potassium channels ^8^. Our results also demonstrate that these channels are widely expressed in cervical spinal motoneurons, including those innervating the diaphragm. A significant fraction of these motoneurons is excited by fentanyl, concomitant with blockade of voltage-dependent non-inactivating K^+^ currents*. In vivo* electromyography revealed a persistent tonic component of diaphragmatic muscle activity elicited by fentanyl, but not morphine. Taken together our results identify a novel off-target mechanism for fentanyl action, independent of μ-opioid receptor activation, with a paradoxical excitatory effect that may underlie WCS. We anticipate these findings may inform the design of safer analgesics and generalize to other neuronal circuits implicated in fentanyl-related maladaptive behaviors.

## Main

The opioid epidemic remains a significant public health threat despite a recent reversal of reported overdose fatalities, driven primarily by the easy availability and the unpredictable variability of illicit doses of fentanyl and fentanyl derivatives ^1^. Rapid death is suggested in ∼50% of overdose victims by post-mortem analysis of plasma levels which revealed extraordinarily high levels of fentanyl, yet negligible levels of its main metabolite norfentanyl ^9^. Similarly, at a supervised injection site in Vancouver, Canada muscle rigidity accompanied ∼48% of overdose cases characterized by atypical overdose presentation, suggesting a link to fatality ^10^. Fentanyl is a highly potent synthetic opioid initially synthesized by Paul Janssen as an analgesic alternative to morphine, created by modifying meperidine to increase its lipophilicity ^2^. Fentanyl and its derivatives are structurally divergent from morphine and its competitive antagonist naloxone, yet both classes of compounds bind to overlapping pockets in the μ-opioid receptor ^11^. Although highly effective as an analgesic, fentanyl when delivered rapidly and at high concentrations can trigger a potentially lethal side-effect termed Wooden Chest Syndrome (WCS) ^6,7^. This results from generalized muscle rigidity, notably of musculature critical for breathing, including jaw and laryngeal muscles that control upper airway patency, and intercoastal and diaphragm muscles required for mechanically inflating the lungs ^5,12,13^. When recognized perioperatively, WCS is treated with competitive blockers of the nicotinic acetylcholine receptor, such as rocuronium, to reduce excessive muscle contraction by dampening neuromuscular transmission ^14,15^. Reports of reversal of WCS by naloxone are inconsistent. Failures indicate that while naloxone may recover opioid-mediated suppression of central respiratory drive, this can often be insufficient for reversing WCS ^16,17^. Together this suggests that the mechanism responsible for WCS lies centrally, downstream of the pattern-generating circuitry for respiratory motor output, but upstream of the effector musculature, and may involve molecular targets other than the μ-opioid receptor. Indeed, several hypotheses have been advanced suggesting a role for disinhibition of local inhibitory spinal premotor circuitry by opioid-sensitive medullary-spinal projections of noradrenergic or serotonergic neurons ^18–20^, or alternatively by direct binding of fentanyl to catecholaminergic receptors ^21,22^. However, presently no strong consensus exists.

Potassium channels of the Ether-à-Go-Go (EAG) superfamily ^8,23–25^ are an attractive candidate for off-target blockade by fentanyl, due to their widespread distribution of expression in the nervous system ^26,27^ and the well-demonstrated propensity for members of this superfamily, particularly HERG (Kv11.1/*KCNH2*), to promiscuously bind and be blocked by many diverse compounds ^28–30^ Recently, fentanyl was reported to reversibly block Kv11.1 by the Zhang lab, with an IC50 of 0.9 μM, while norfentanyl, the main metabolite of fentanyl, exhibited a ∼100-fold lower affinity and naloxone failed to block entirely ^31^. We hypothesized that WCS may result in part from fentanyl-induced excitation of respiratory motoneurons due to the direct block of EAG-class potassium channels expressed in these motoneurons, leading to increased intrinsic excitability, independent of signaling through the μ-opioid receptor. In turn, this may result in excessive stimulation of respiratory muscles and uncontrolled tonic contractions.

### Fentanyl blocks *Eag/Erg* K^+^ channels

To test this hypothesis, we demonstrated that fentanyl directly blocks a subset of EAG-class potassium channel sharing the closest amino acid sequence similarity to the residues of HERG (Kv11.1/*KCNH2*) that form the ion conduction pathway. The EAG superfamily of potassium channels comprises of eight genes in mammals which can be subdivided into three subclasses based on sequence similarity. These are termed *Eag* (Kv10.1/*KCNH1*, Kv10.2/*KCNH5*), *Erg* (Kv11.1/*KCNH2*/HERG, Kv11.2/*KCNH6*, Kv11.3/*KCNH7*) and *Elk* (Kv12.1/*KCNH8*, Kv12.2/*KCNH3*, Kv12.3/*KCNH4*) (Figs. 2a, S3) ^8,32,33^. We focused on the *Eag*- and *Erg*-subclasses of potassium channels because they exhibit the highest conservation of the three critical residues in HERG (S624, Y652 and F656) previously identified by the Sanguinetti lab, to be required for binding of MK-499, a HERG-specific blocker, by alanine-scanning through the ion conduction pathway ^34^. Five expression plasmids encoding *Eag* (Kv10.1, Kv10.2) ^35,36^ and *Erg* (Kv11.1, Kv11.2, Kv11.3) ^23,24,34^ subclasses of potassium channels were expressed in HEK293 cells by acute transfections, followed by patch-clamp recordings. All the constructs, except Kv11.2, yielded sufficient currents matching those previously reported, for subsequent assays (Figs. 1a, S1). Fentanyl (10 μM) reversibly blocked all four expressed *Eag* and *Erg* currents, in a voltage-dependent and subtype-specific manner. The two *Erg* potassium channels (Kv11.1, Kv11.3) exhibited the greatest block of ∼70% maximal conductance (Gmax) (p= 0.0022 and p= 0.0006 respectively; unpaired Mann-Whitney), whereas the *Eag*-subclass Kv10.1 potassium channel exhibited ∼50% block of Gmax (p= 0.0022, unpaired Mann-Whitney), and Kv10.2 showed the weakest but significant block of ∼20% Gmax (p= 0.0022, unpaired Mann-Whitney). Both blocking action and partial reversal by washout were rapid, within the ∼30 secs required for the exchange of our bath volume, consistent with a direct reversible block of these channels (Kv10.1, 80%, p= 0.0033; Kv10.2, 90%, p= 0.0010; Kv11.1, 110%, p= 0.0075; Kv11.3, 75%, p= 0.0003; paired t-test). These measurements were unaffected by potential rundown under our recording conditions, with the concentration of Ca^2+^ in our internal pipette solution highly buffered to ∼9 nM ^37^. Interestingly, we observed a significant rebound of HERG current with washout, similar to that previously reported ^31^, suggesting that fentanyl may bind with a low and high affinity site.

**Fig 1.**
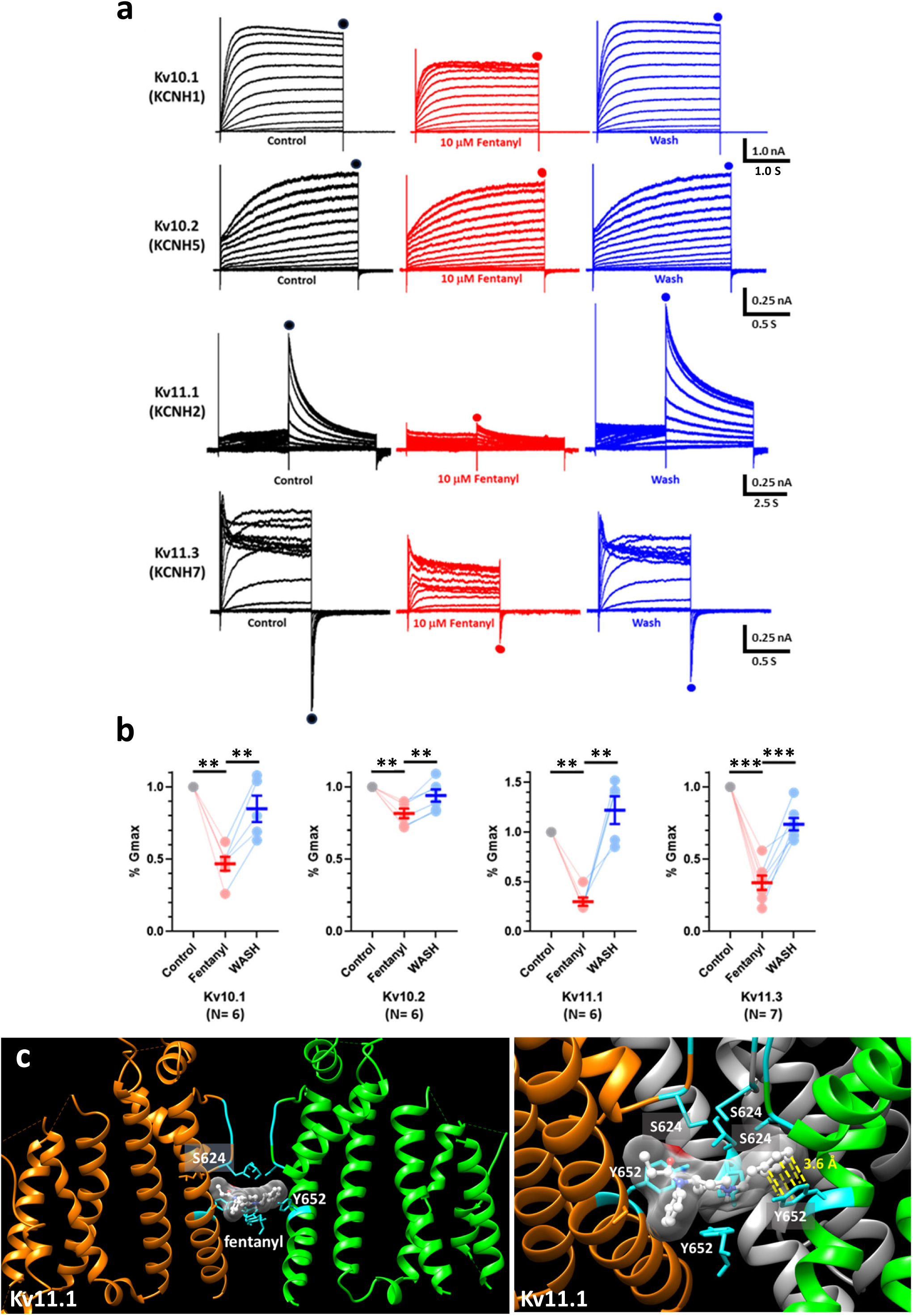
*Eag-* and *Erg*-subtypes of EAG K^+^ channels are reversibly blocked by fentanyl. **a.** Kv10.1(*KCNH1*), Kv10.2(*KCNH5*), Kv11.1 (*KCNH2*) and Kv11.3(*KCNH7*) currents expressed in HEK293 cells. Peak current used for calculating conductance (G), measured as isochronic currents at the end of voltage steps for Kv10.1 and Kv10.2, and as inward tail currents for Kv11.3. For Kv11.1, due to rapid inactivation, peak currents measured as outward tail currents at −50 mV due to rapid recovery from inactivation and much slower deactivation kinetics (see Methods for details of voltage protocols and Fig 1S for G/V plots). **b.** Bath applied fentanyl (10 μM) reversibly blocks maximal conductance (Gmax) for all constructs, in a *Eag/Erg*-subtype specific fashion: Kv10.1 (50%; p= 0.0022), Kv10.2 (20%; p= 0.0022), Kv11.1 (75%; p= 0.0022), and Kv11.3 (70%; p=0.0006). Block was partially reversed by wash: Kv10.1 (80%; p= 0.0033), Kv10.2 (90%; p= 0.0010), Kv11.1 (110%; p= 0.0075), Kv11.3 (75%; p= 0.0003). Statistical significance for block determined by unpaired Mann-Whitney comparisons between control and fentanyl values, and for reversal by paired t-test comparisons between fentanyl and wash values. **c.** Molecular prediction of the fentanyl binding site in Kv11.1 by Autodock-Vina, within the rings defined by S624 at the base of the pore α-helix and Y652 at the top of S6 in each of the four channel subunits (left panel). Fentanyl docks asymmetrically relative to the central axis of the ionic conduction vestibule (also see Fig. S2c) and exhibits a close 3.6 Å parallel orientation of its phenethyl ring to the phenolic ring of Y652, predictive of a strong stacked π-orbital molecular interaction (right panel). Ribbon representations of only two subunits in the left panel and three subunits in the right panel displayed for clarity.

**Fig. 2.**
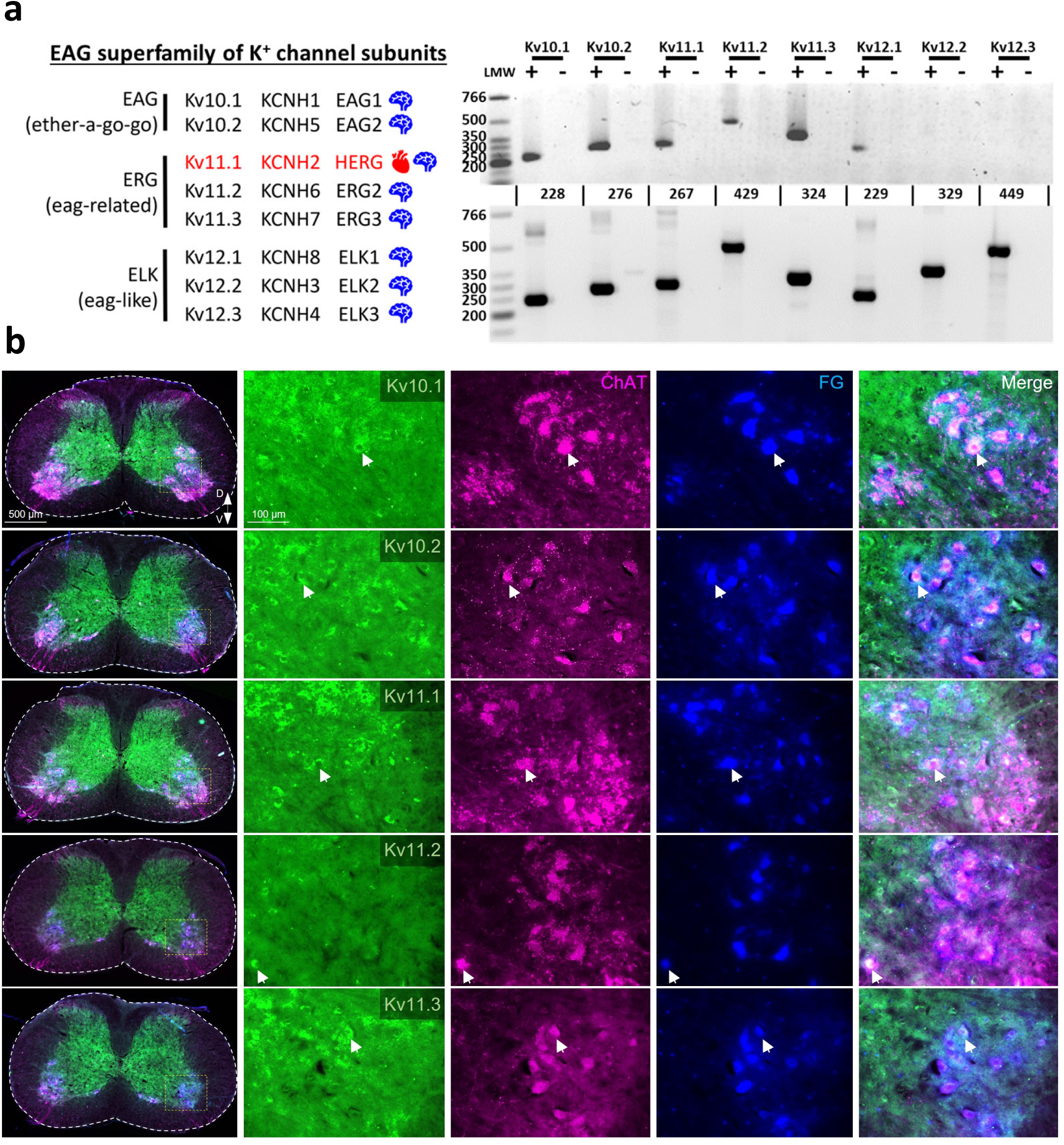
Selective expression of transcripts encoding *Eag*/*Erg*-subtypes of EAG K^+^ channel subunits in phrenic motoneurons. **a.** RT-PCR assays with total RNAs from micro-dissected spinal C3-5 ventral horn tissue and whole brain with primer pairs specific to each of the 8 genes encoding all EAG subtypes of K^+^ channel subunits. Selective enrichment for the expression of only *Eag* (Kv10.1, Kv10.2)- and *Erg* (Kv11.1, Kv11.2, Kv11.3)-subtypes, with little to no expression detected for *Elk* (Kv12.1, Kv12.2, Kv12.3)-subtype transcripts. By contrast, all EAG transcripts were readily detected from whole brain total RNA. Parallel control assays with first-strand reactions performed without reverse transcriptase (-) were negative. All assays repeated 4X with similar results. **b.** Detection of individual *Eag/Erg* transcripts in single phrenic motoneurons by *in situ* hybridization with RNAscope probes. Phrenic motoneuron in spinal cervical segments C3-5 were identified by retrograde labeling by injections of fluorogold (FG) in the lateral diaphragm, and immunolabeling for choline acetyltransferase (ChAT). Shown are examples of phrenic motoneurons (FG^+^, ChAT^+^), co-labeled with RNAscope probes specific to each of the *Eag/Erg*-subtype transcripts. Quantitation revealed nearly all phrenic motoneurons expressed *Eag* transcripts [Kv10.1 (∼96%) and Kv10.2 (85%)], and ∼60% expressed two of the three *Erg* transcripts [Kv11.1 (66%), Kv11.3 (60%), Kv11.2 (22%)] (see Methods for details of quantitation).

To gain insight into the potential structure of the binding sites for fentanyl in these *Eag* and *Erg* potassium channels, we used AutoDock-Vina ^38,39^ to computational dock fentanyl into the available structures of Kv10.1 (8EOW), Kv10.2 (7YIH) and Kv11.1 (8ZYO) derived by single particle cryogenic electron microscopy (cryo-EM) ^40–42^. Using a search volume of 20-25 Ǻ^3^ centered around the central conduction pathway, fentanyl was predicted to dock in all three channel structures. Fig. 1c illustrates the predicted docked structure of the HERG-fentanyl complex, showing fentanyl binding at the top of the conduction pathway, in the space defined by two rings. These rings are formed by S624 at the base of the pore α-helix and Y652 near the top of S6, previously identified to be critical for MK-499 binding. In addition, the benzene ring of the phenethyl moiety of fentanyl aligned in parallel with the phenolic ring of tyrosine at Y652, to allow a close interaction by π-orbital stacking, predictive of a strong molecular interaction. As with the Kv11.1-astemizole structure ^41^, the docked fentanyl binding site is asymmetric relative to the central axis of the conduction pathway. However, the electrostatically positive central piperidine ring of fentanyl is sufficiently lodged to replace and occlude K^+^ ions that would otherwise be focally attracted to the base of the selectivity filter, due to a region of high electronegativity created by dipolar charge separation at the C-terminal ends of the four pore α-helices ^43^ (Fig. S2c).

The predicted Kv10.1-fentanyl docked structure is similar to that for Kv11.1-fentanyl (Fig. S2a). Fentanyl is predicted to bind to Kv10.1 between the two rings formed by S463 and Y491, analogous to Kv11.1. However, the central piperidine ring of fentanyl is positioned more symmetrically, centered directly underneath the selectivity filter. Interestingly for Kv10.2, which exhibited the weakest blocking with fentanyl, AutoDock-Vina was unable to dock fentanyl between the same rings (T432 and Y460) analogous to Kv11.1 (Fig. S2b). Instead, fentanyl was predicted to dock more peripherally within the conduction pathway, below Y460, and in many more conformations, consistent with a lower affinity interaction between the ligand and Kv10.2. Kv11.3 was not explicitly modeled. But, due to complete sequence identity between Kv11.3 and Kv11.1 from the pore α-helix through S6, we anticipate a similar fentanyl docking profile between these two *Erg* potassium channels. These results and predicted interactions between these *Eag*/*Erg* channels and fentanyl may inform the future design of fentanyl derivatives that preserve efficacy at the μ-opioid receptor, without blocking these potassium channels.

### *Eag/Erg* express in spinal motoneurons

To examine the expression pattern of *Eag* and *Erg* subclasses of potassium channels, RT-PCR assays were performed with gene-specific primers for all *Eag* and *Erg* subclasses of potassium channels using total RNAs isolated from micro-dissected ventral horn tissue from spinal cervical segments 3-5 slices containing phrenic motoneurons that innervate the diaphragm, and from the whole brains of adult mice. These RT-PCRs revealed the expression of all *Eag* (Kv10.1. Kv10.2) and *Erg* (Kv11.1, Kv11.2, Kv11.3) transcripts in C3-5 ventral horn tissue, but little to no expression of *Elk* (Kv12.1, Kv12.2, Kv12.3) transcripts. In contrast, the expression of all EAG channel subunits were readily detected in whole brain RNA samples, indicating the efficacy of all our primer pairs (Fig. 2b).

To examine the expression pattern of *Eag* and *Erg* transcripts at greater cellular resolution, *in situ* hybridizations were performed with RNAscope probes specific to each *Eag*/*Erg* gene. Spinal C3-5 serial sections were prepared from mice injected with fluorogold (FG) in the diaphragm to retrogradely label phrenic motoneurons. These spinal sections were hybridized with RNAscope probes either singly or in combinations, and counter-labeled for choline acetyltransferase (ChAT) by immunostaining. This allowed us to localize specific expression of individual transcripts in single phrenic motoneurons, as well as the coexpression of pairs of *Eag*/*Erg* transcripts in individual motoneurons. We found widespread expression of all *Eag* and *Erg* transcripts in a significant fraction of all FG-labeled phrenic motoneurons, and in unlabeled motoneurons based on the morphological criteria of location and large soma size, and ChAT^+^ signals. Counts of RNAscope signals over phrenic motoneurons identified by both FG and ChAT labelling revealed that nearly all phrenic motoneurons expressed *Eag* transcripts (Kv10.1: 96±2%, Kv10.2: 85±7%), and roughly 60% expressed two of the three *Erg* transcripts (Kv11.1: 66±12%, Kv11.2: 22±5%, Kv11.3: 60±13%). We also found evidence for the co-expression of multiple pairs of *Eag*/*Erg* transcripts ^44^ in individual motoneurons, including Kv10.1 + Kv10.2, Kv11.1 + Kv10.1 and Kv11.1 + Kv10.2. We conclude that a significant fraction of C3-5 motoneurons, including phrenic motoneurons which innervate the diaphragm, express a variable palette of *Eag* and *Erg* potassium channel subunits, presumably to tune the electrical properties of individual motoneuron.

### Fentanyl excites phrenic motoneurons

To directly examine the impact of acutely applied fentanyl on the electrical properties of phrenic motoneurons, whole-cell patch-clamp recordings were performed on *in vitro* mouse slices of C3-5 spinal cords, targeting motoneurons either identified based on morphological criteria, or in some cases by prior retrograde labeling with Alexa594-conjugated cholera toxin B (CTB) injected in the diaphragm to visually identify phrenic motoneurons. Electrical properties were measured under current-clamp conditions in response to progressive current injections, or under voltage-clamp conditions in response to voltage-step protocols. In addition, all recordings were performed under pharmacological blockers for synaptic transmission against ionotrophic receptors for glutamate, GABA, and glycine (20 µM CNQX, 10 µM CPP, 1 µM gabazine, and 1 µM strychnine, respectively), to isolate intrinsic membrane properties. Similar results were observed for unlabeled motoneurons and phrenic motoneurons identified by CTB labeling (Fig. 3a-b).

**Fig. 3.**
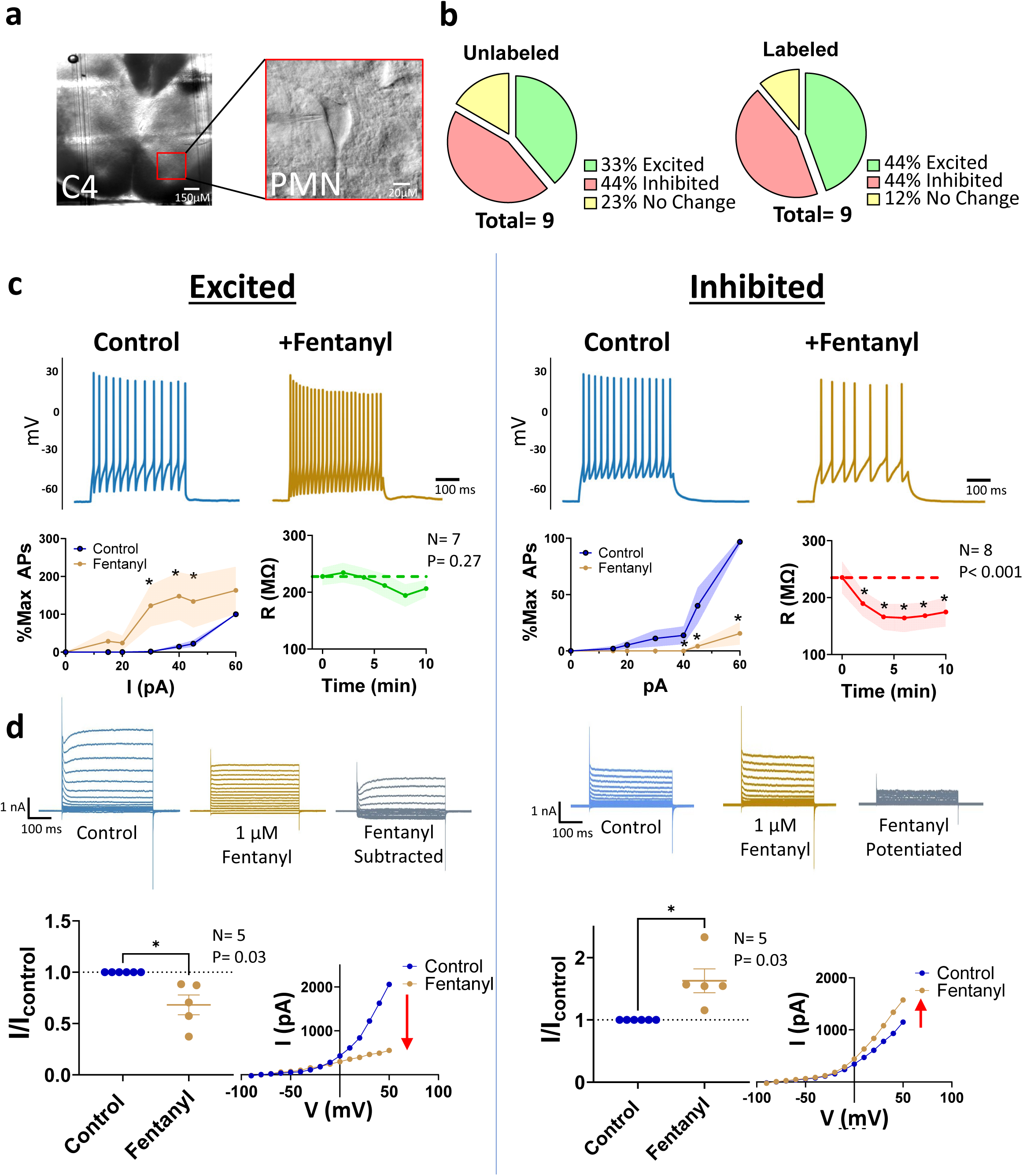
Fentanyl differentially excites and inhibits subpopulations of spinal C3–5 phrenic motoneurons. **a.** Representative transverse section from the C4 spinal cord segment illustrating the region targeted for motoneuron recordings (left panel), along with an example of a targeted phrenic motoneuron identified through retrograde labeling with Alexa 594-conjugated cholera toxin B (CTB) or morphological characterization (right panel). **b.** Approximately 44-33% of neurons, identified via retrograde CTB labeling or morphological criteria, exhibited excitatory responses following bath application of 1 µM fentanyl. In contrast, 44% of neurons were inhibited, and ∼12–23% displayed no response to fentanyl application. **c.** Spinal motoneurons excited by fentanyl demonstrated significantly elevated spike rates during stepwise current injections, with minimal changes in input resistance (R), compared to pre-fentanyl control conditions (left panel). Conversely, motoneurons inhibited by fentanyl exhibited a significant reduction in spike rates and input resistance during identical current injection protocols (right panel). **d.** Under voltage-clamp, fentanyl-excited motoneurons showed a significant reduction in isolated outward currents during voltage-step protocols following fentanyl application. In contrast, fentanyl-inhibited motoneurons displayed an increase in isolated outward currents under the same conditions. Representative current-voltage (IV) curves are shown for motoneurons excited or inhibited by fentanyl.

Current-clamp recordings revealed that a significant population (∼44%) of phrenic motoneurons exhibited a rapid increase in excitability after bath applied fentanyl (1 μM), which was reflected in a significant increase in the frequency of action potentials elicited by current injections over baseline controls with 30, 40, and 45 pA current injections (p= 0.03, N= 7, two-way RM ANOVA). In many cases, fentanyl application was accompanied by a slight depolarization of resting membrane potential of 5-10 mVs, although this was not significantly reflected by a corresponding change in resting input resistances (Fig. 3c). These results are consistent with fentanyl decreasing the threshold for triggering action potentials in this pool of motoneurons, by altering a voltage-dependent conductance. By contrast, ∼44% of phrenic motoneurons were electrically inhibited by fentanyl under similar current-clamp conditions, resulting in a significant decrease in the frequency of action potentials elicited by current injections of 45 and 60 pA (p<0.001, N= 8, two-way RM ANOVA). In this pool of motoneurons, decreases in excitability were accompanied by a significant 50% decrease in resting input resistance (p< 0.001, N= 8, one-way RM ANOVA). This decreased excitability is consistent with an altered conductance active at resting membrane potentials. The remaining pool of ∼12% of phrenic motoneurons exhibited no detectable effect on excitability with fentanyl.

To specifically examine K^+^ currents, voltage-clamp recordings were performed in the presence of TTX (1 μM) and Cd^2+^ (1 mM) to block voltage-gated sodium and calcium channels, respectively, in addition to the same panel of pharmacological blockers of synaptic transmission as mentioned above (Fig. 3d). In the pool of motoneurons excited by fentanyl, a non-inactivating outward voltage-gated K^+^ current was observed under control conditions, which was reduced by 32±10% (p= 0.03, N= 6, paired t-test) in the presence of fentanyl (1 μM). Subtraction of fentanyl-blocked currents from controls, revealed a large component of initial outward currents resembling a voltage-gated K^+^ current with “delayed rectifier”-like properties, similar to some heterologously expressed *Eag*/*Erg* currents ^26^. By contrast, in motoneurons inhibited by fentanyl, voltage-clamp recordings revealed a 62±27% increase in potassium conductance (p= 0.03, N= 6, paired t-test), not gated by voltage. Subtractions of control from fentanyl-treated currents, revealed a residual fentanyl blocked current with near instantaneous kinetics and no intrinsic voltage-dependent gating. This residual component of fentanyl-activated potassium current is consistent with inward rectifier GIRK K^+^ currents, known to be activated by agonist-bound μ-opioid receptors.

Taken together, these recordings from C3/5 spinal motoneurons *in vitro* provide evidence that a significant fraction (∼40%) of motoneurons, including those that innervate the diaphragm, are electrically excited by fentanyl by a direct block of a distinctive outward K^+^ current. Although these currents resemble those of some *Eag*/*Erg* channels ^26^, we cannot exclude the possibility that other yet to be identified classes of potassium channels with “delayed rectifier”-like properties may also contribute to this component of fentanyl-blocked outward current. Our data also shows that the electric response to fentanyl in motoneurons is heterogenous, with a large fraction of motoneurons exhibiting inhibition by fentanyl, likely contributed to by the activation of GIRK conductance through a conventional μ-opioid receptor Gα_i/o_ signaling pathway ^3,4^. This heterogeneity may reflect the functional diversity of phrenic motoneurons, including those that adapt rapidly to provide phasic neuromuscular stimulation, while other are relatively non-adapting and recruited later during the inspiratory phase to provide tonic stimulation to the diaphragm ^45,46^. The *in vivo* consequences of fentanyl on respiratory muscle contractions are thus likely to be determined by a summation of its different effects on multiple pools of motoneurons in the whole organism. Moreover, the outcome of these concurrent inhibitory and excitatory actions will likely be state-dependent which may explain why the presence and degree of WCS is unpredictable, making this phenomenon particularly dangerous.

### Fentanyl induces tonic muscle activity

To examine the *in vivo* effect of fentanyl delivered at high doses (500 μg/kg, ip) on diaphragm activity, electromyogram (EMG) recordings were performed on urethane-anesthetized adult mice in response to either fentanyl or morphine (Fig. 4) ^47^. Fentanyl delivered by intraperitoneal injection caused a rapid reduction of respiratory rate from ∼240 breaths/min to a stabilized frequency of ∼110 breaths/min after 15 minutes. Control animals injected with morphine exhibited the same reduction of respiratory rate with a similar time course, using a dose calibrated to suppress respiratory drive to the same extent (150 mg/kg, ip). However, only fentanyl-injected animals also evoked a tonic component of integrated EMG activity, corresponding at its peak to ∼20-40% of the maximal integrated EMG activity of pre-fentanyl control EMGs. Inspection of the raw EMG recordings revealed that this tonic component is the result of the recruitment of tonic motor units. These motor units fire between phasic inspiratory bursts, inducing an increased constitutive tone of diaphragm contraction throughout the respiratory cycle, indicative of WCS. Morphine-injected control animals failed to exhibit a similar increase in diaphragmatic muscle tone. Interestingly, this fentanyl-induced WCS effect was transient. Being rapidly induced, motor activity returned to near baseline control levels within 15 mins. By contrast, suppression of central respiratory drive, as reflected by the diminished frequency of the phasic component of EMGs, persisted beyond the time of recovery of the WCS effect. These different time courses for the effect of fentanyl are consistent with its relatively rapid pharmacokinetic properties. As previously described, free levels of circulating fentanyl are lowered by partitioning to lipid deposits and metabolism to norfentanyl in the liver ^48^. The blockade of *Eag*/*Erg* potassium channels by fentanyl is achieved by relatively low affinity binding, compared to its high binding affinity for activation of μ-opioid receptors. Thus, we hypothesize that the WCS-effect on motoneurons recovers before central respiratory drive, as initial high levels of circulating fentanyl levels are lowered by *in vivo* partitioning and metabolic catabolism, relieving *Eag/Erg* potassium channel blockade but sustaining μ-opioid receptor activation.

**Fig. 4.**
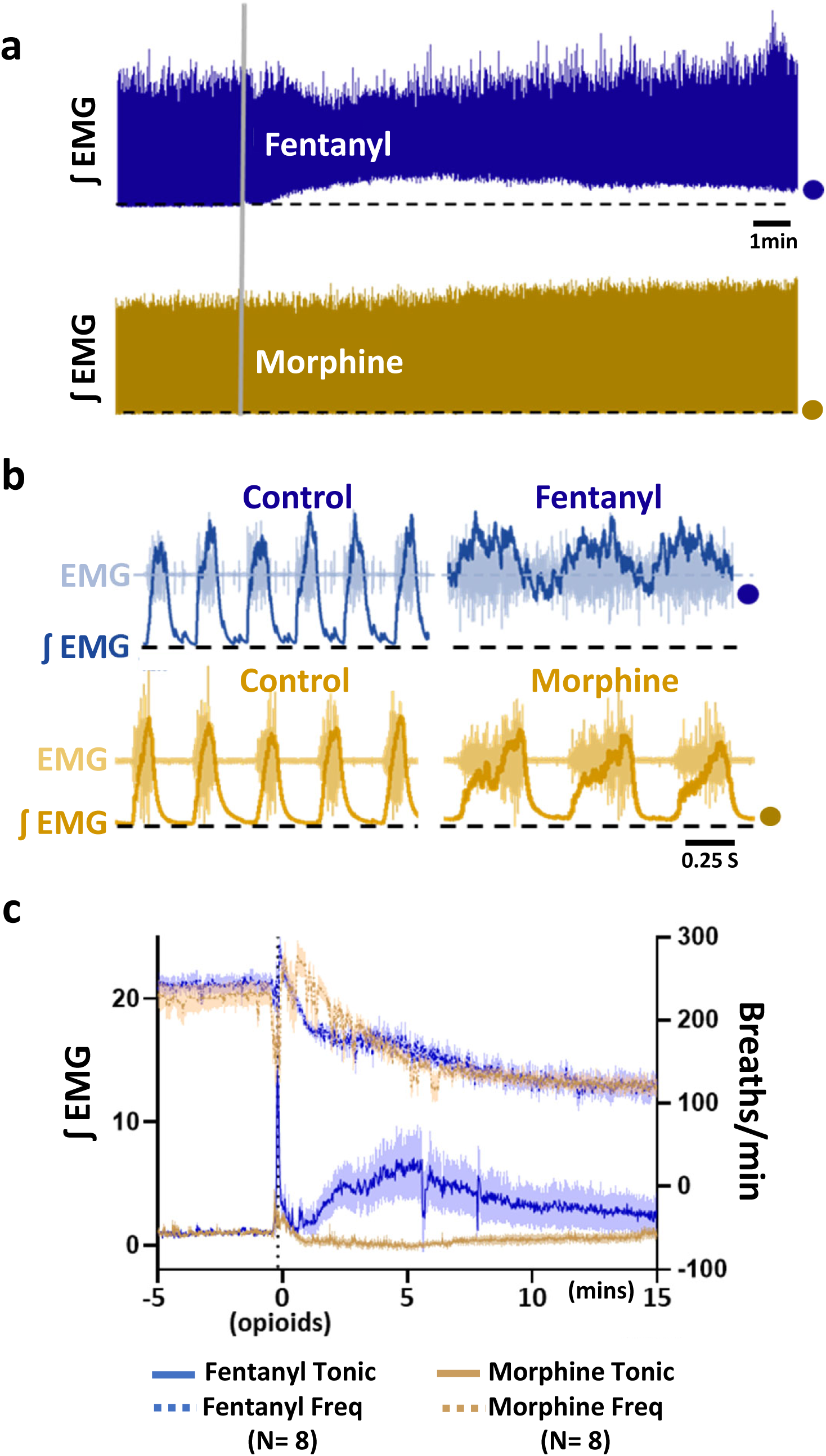
Fentanyl uniquely induces tonic activation of the diaphragm. **a.** Intraperitoneal (IP) administration of fentanyl (500 µg/kg) resulted in a sustained increase in baseline activity of the integrated diaphragmatic EMG, characterized by tonic activation throughout the respiratory cycle. This tonic activity persisted during the expiratory period, preventing full relaxation of the diaphragm. This phenomenon was unique to fentanyl and not observed following morphine administration (150 mg/kg IP), despite both opioids causing similar respiratory suppression. **b.** Expanded raw and integrated diaphragmatic EMG recordings highlight the fentanyl-induced tonic activation, which contrasts with the lack of tonic activity observed following morphine treatment. **c.** The fentanyl-induced tonic activation of diaphragm activity (solid blue line) emerged rapidly following administration, peaking approximately 5 mins post-injection. This effect was not observed with morphine administration (solid gold line), even though both fentanyl and morphine caused comparable levels of respiratory suppression, as indicated by decreased respiratory rates (dotted blue and gold lines, respectively).

In summary, we identified a subclass of EAG potassium channels that is reversibly blocked by fentanyl and provide a predicted molecular characterization of their fentanyl binding sites. We demonstrate that the genes encoding these EAG potassium channels are expressed in spinal cervical C3-5 motoneurons, including phrenic motoneurons that innervate the diaphragm.

Recordings from these motoneurons revealed a significant fraction (∼40%) which are electrically excited by fentanyl, along with additional pools of motoneurons that are inhibited or unaffected. This excitation was concomitant with the blockade of a voltage-gated outward K^+^ current with “delayed rectifier”-like properties similar to some EAG potassium channels ^26^. *In vivo*, high doses of fentanyl recapitulated the hallmarks of WCS in diaphragm muscle, by eliciting a tonic component of muscle activity, consistent with recruitment of tonic neuromuscular stimulation evoked by fentanyl. These results provide support for a novel mechanism of fentanyl action, based on off-target blockade of a subset of EAG channels, resulting in paradoxical excitation. Future studies will examine if other classes of potassium channel contribute to WCS by the same off-target blocking mechanism. *Eag*/*Erg* potassium channels are widely expressed in the CNS, including in the DA2 cluster of dopaminergic neurons in the ventral tegmental area implicated in addictive behaviors (Fig. S6) ^49^. We speculate that this same blocking mechanism demonstrated for WCS may generalize in unsuspected ways to other fentanyl-related behaviors such as addiction and drug seeking^50^, mediated by supra-medullary neuronal circuits unrelated to respiratory control.

## Supporting information

Suppl-Wei et al-WCS

## Materials and Methods

### Ethical Approval

Mouse experiments were performed on male and female adult (P50-80) and juvenile (P4-12) C57BL/6J mice bred at Seattle Children’s Research Institute in accordance with Seattle Children’s Research Institute Animal Care and Use Committee (Protocol IACUC00058) and National Institutes of Health guidelines. Adult mice weighed between 28-30 g and both adult and juvenile mice were group housed and given access to food and water *ad libitum*. Light/dark cycles were maintained at 12 hr each and temperature controlled at 22±1 °C.

### HEK293 Electrophysiology

HEK293 cells (CRL-1573; ATCC, Gaithersburg, MD) were cultured in standard media [DMEM, high glucose, GlutaMAX (Gibco 10566016; Thermo Fisher Scientific, Waltham, MA), supplemented with 10% (v/v) fetal calf serum (FCS) (Gibco A5670401; Thermo Fisher Scientific, Waltham, MA) and 1% (v/v) Penicillin-Streptomycin (P/S) (10,000 U/ml) (Gibco 15140148; Thermo Fisher Scientific, Waltham, MA)] at 37°C and 5% CO_2_. Cells were passaged in T25 tissue culture flasks (FB012935; Thermo Fisher Scientific, Waltham, MA) approximately once a week. Only cells passaged < 20X were used for expression studies.

Plasmid constructs were acutely transfected into HEK293 cells using Viafect reagent (E4981; Promega, Madison, WI), following the manufacture’s protocol. In brief, HEK293 cells were prepared for transfection by plating into 12-well tissue culture plates (Nunc 12-565-321; Thermo Fisher Scientific, Waltham, MA) at a density of ∼0.5-2 ×10^5^ cells per well and grown to ∼80-90% confluence, allowing for one confluent well per transfection condition. On the day of transfection, media in wells to be transfected were replaced with 0.5 ml fresh DMEM with 10% FCS, without P/S. Lipophilic/DNA transfection complexes were generated for each well to be transfected by combining ∼1.0 μg of plasmid DNAs with serum-free OptiMEM (Gibco31985062; Thermo Fisher Scientific, Waltham, MA) to a final volume of 100 μL, then adding 3.0 μL Viafect and allowing the mixture to assemble for 30 mins at 24°C. Assembled transfection mixtures were then added to wells, dropwise. Transfected cells were incubated overnight at 37°C, and visually monitored for transfection efficiency *in situ* using a plate microscope equipped with fluorescence (Invitrogen EVOS M7000; Thermo Fisher Scientific, Waltham, MA). Transfection efficiencies were typically >70%.

The following *Eag/Erg* expression plasmids were used:

1. KCNH1/Kv10.1 (human); pcKv10.1, a gift from Luis Pardo, MPI-Gottingen, Germany (Addgene 85703).
2. KCNH5/Kv10.2 (human); pTKv10.2, a gift from Luis Pardo, MPI-Gottingen, Germany (Addgene 85706).
3. KCNH2/Kv11.1 (human); pCEP4-herg1a, a gift from Michael Sanguinetti, University of Utah, USA (Addgene 53052).
4. KCNH6/Kv11.2 (rat); rerg2-pcDNA3, a gift from Christiane Bauer and Robert Bahring, University Medical Center-Hamburg/Eppendorf, Germany.
5. KCNH7/Kv11.3 (rat); rerg3-pcDNA3, a gift from Christiane Bauer and Robert Bahring, University Medical Center-Hamburg/Eppendorf, Germany.

As a visual marker for transfections, 50-100 ng of pcDNA3 expressing GFP was added to each lipophilic/pDNA transfection mixture.

Following overnight incubation, transfected cells were dissociated with TrypLE (Gibco 12-604-013, Thermo Fisher Scientific, Waltham, MA), and replated at low density onto poly-D-lysine-coated 12 mm diameter glass coverslips (GG-12-pdl; NeuVitro, Vancouver, WA) in 24-well tissue culture plates (FisherBrand FB012929; Thermo Fisher Scientific, Waltham, MA), for patch-clamp electrophysiology. Typically, ∼10,000-15,000 cells were replated per well allowing sufficiently low density to isolated individual cells. This prevented formation of electrical junctions between contacting cells which would hinder adequate space-clamp recording conditions. Recordings were performed 0.5-2 days after replating at 24°C.

For patch-clamp recordings, coverslips containing adherent cells were transferred to the stage of a Zeiss AxoExaminer.A1 microscope, equipped with a 40X water immersion objective and epifluorescence optics. Pipettes were positioned with a Sutter MPC-325 micromanipulator (Novato, CA). Whole-cell voltage-clamp recordings were acquired with an AxoClamp200B amplifier (Molecular Devices, Union City, CA), using pClamp10.4. The compositions of recording solutions were as follows: Bath (in mM, 4.0 KCl, 145 NaCl, 2.0 CaCl_2_, 2.0 MgCl_2_, 10 glucose, 10 HEPES, pH 7.4); Pipette internal solution (in mM, 140 K-gluconate, 4.0 Na_2_ATP, 1.0 CaCl_2_, 2 MgCl_2_, 10 EGTA, 10 HEPES, pH 7.2). Under these conditions, the [Ca^2+^] of the pipette internal solution was buffered to ∼9 nM to minimize rundown of *Eag/Erg* currents, previously shown to be due to binding of Ca^2+^/Calmodulin to the N-terminus of these channel subunits ^1^. Patch pipettes were pulled from borosilicate glass (1B120F-4; World Precision Instruments, Sarasota, FL) on a P97 Sutter Instruments puller (Novato, CA) with tip resistances of 3-4 MΩ. Currents were allowed ∼3-5 mins to stabilize after achieving whole-cell recording configuration and acquired at 10 kHz and filtered at 5 kHz. Series resistance compensation was >90% for all recordings. Drug applications were performed by gravity-fed exchange of bath volumes corresponding to ∼2-3X bath volume (0.9 mL), over 30 sec. Fentanyl (#07-890-5657) and morphine (#07-892-4699) stock solutions were obtained from Patterson Vet.

Voltage protocols used for individual *Eag/Erg* construct were as follows:

1. Kv10.1/*KCNH1*. From a holding potential of −80 mV, 2.0 sec voltage steps were taken from −90 to 70 mV in increments of 10 mV, returning to −80 mV holding potential, with a 6 sec interpulse interval.
2. Kv10.2/*KCNH5*. From a holding potential of −80 mV, 1.5 sec voltage steps were taken from −90 to 70 mV in increments of 10 mV, returning to −80 mV holding potential, with a 6 sec interpulse interval.
3. Kv11.1/*KCNH2*. From a holding potential of −80 mV, 4.0 sec voltage steps were taken from −90 to 70 mV in increments of 10 mV, then stepped to −50 mV for 5 sec to measure outward tail currents, before returning to −80 mV holding potential. Voltage steps were repeated with a 16 sec interpulse interval.
4. Kv11.3/*KCNH7*. From a holding potential of −100 mV, 1.0 sec voltage steps were taken from −100 to 60 mV in increments of 10 mV, before returning to −100 mV for 2 sec to measure inward tail currents. Voltage steps were repeated with a 6 sec interpulse interval.

No reliable expression of Kv11.2/*KCNH6* was observed, sufficient for subsequent drug assays.

For calculations of conductance, peak currents were measured as isochronic currents at the end of voltage steps for Kv10.1 and Kv10.2, or as instantaneous inward tail currents for Kv11.3. For Kv11.1, due to rapid inactivation characteristic of this potassium channel, peak currents were measured as maximal currents carried by outward tail currents at −50 mV, due to the combination of rapid voltage-dependent recovery from inactivation and slower deactivation kinetics ^2^. For conductance/voltage (G/V) plots, conductance (G) was calculated by the formula:

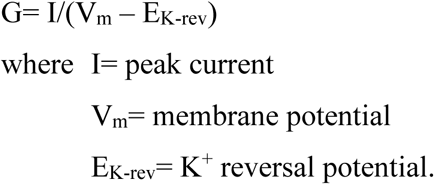

K^+^ reversal potential was −89.6 mV, based on a Nernst equilibrium potential and the recording solutions used.

Current traces were analyzed and plotted using pClamp10.4 (Molecular Devices, San Jose, CA) and Origin 8.5 (Northampton, MA). All G/V curves were plotted as mean with standard error (SE) using Origin 8.5. All statistical calculations were performed in Prism (GraphPad, La Jolla, CA). Figures composed in Microsoft PowerPoint (Redmond, WA).

### Molecular Docking and Visualizations

Molecular docking was performed using AutoDock-Vina ^3^, implemented through the web server SwissDock (www.swissdock.ch) ^4,5^, using coordinates from the cryoEM structures of Kv10.1 (8EOW) ^6^, Kv10.2 (7YIH) ^7^, and Kv11.1 (8ZYO) ^8^, a search volume of 20-25 Å^3^ centered along the central ion conduction pathway, and default parameters. Visualizations were performed with UCSF-Chimera ^9^, developed by the Resource for Biocomputing, Visualization and Informatics at the University of California, San Francisco, with support from the NIH P41-GM103311.

### RT-PCR assays

Total RNA was isolated from whole adult mouse brains or isolated ventral horn tissue samples micro-dissected from spinal cervical C3-5 vibrotome sections, using a Monarch Total RNA miniprep kit (New England Biolabs #T2010S), following the manufacture’s protocol. Pooled ventral horn samples from ∼8 slices (∼25 mg) yielded ∼2.4 μg of total RNA.

Primer pairs for each *Eag/Erg/Elk* gene were designed to target sequential exonic sequences flanking large introns (>2 Kb) to minimize amplification of residual genomic DNA. Primers targeted exons preserved in all splice variants for each gene, based on data in Ensembl (www.ensembl.org). Primer sequences used were:

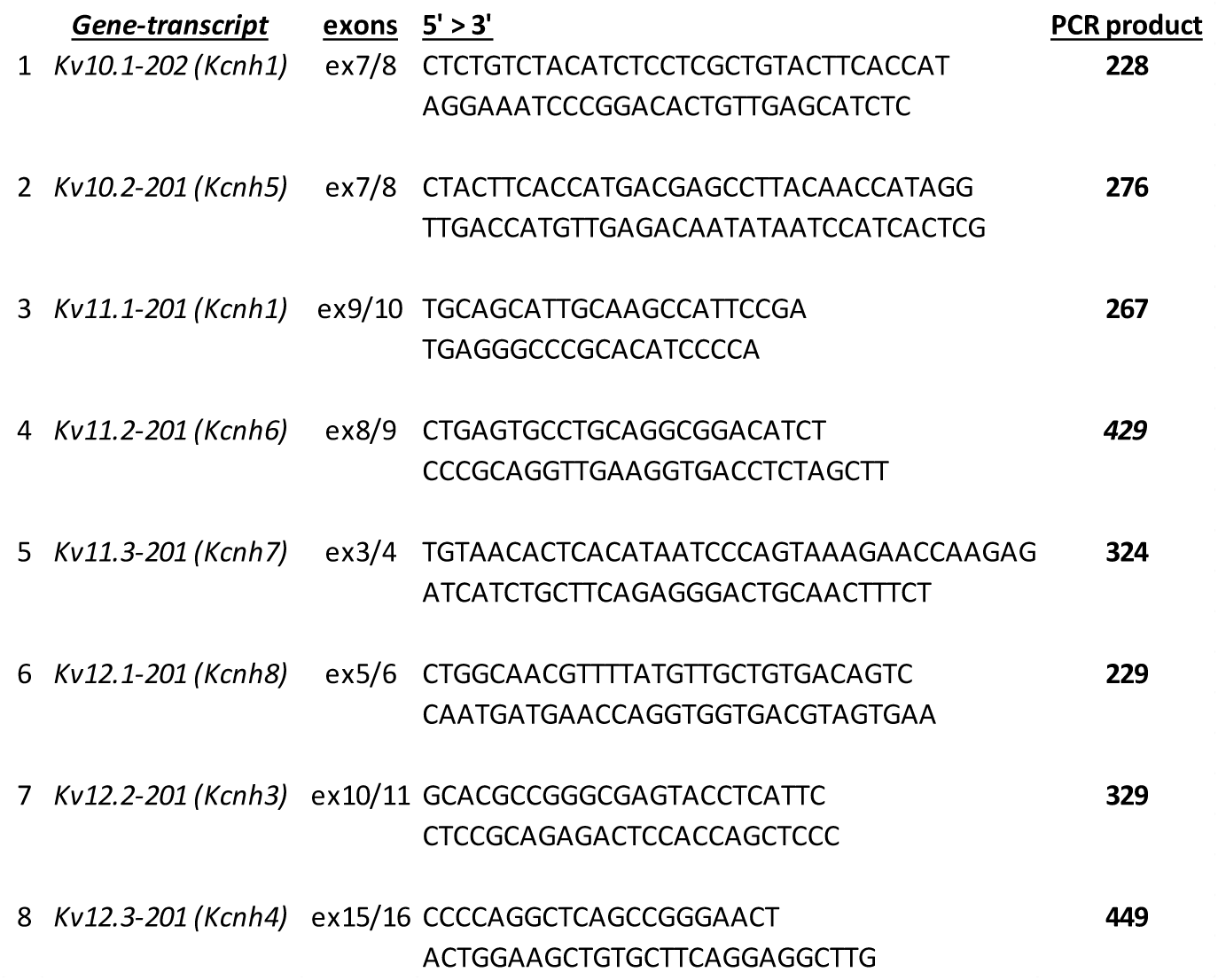

First-strand cDNA synthesis was achieved by standard reactions with M-MuLV reverse transcriptase (NEB #M0253S) using ∼0.25 μg of total RNAs and random hexamer primers, including parallel control reactions without reverse transcriptase. Approximately 1.25% of each first-strand cDNA reaction was used for each PCR assay. PCR assays were performed with Q5-HotStart polymerase (NEB #M0515S) and the following thermocycling protocol:

Step 1: 98°C, 30 sec (polymerase activation)
Step 2: 98°C, 5 sec (denaturing)
Step 3: 68°C, 10 sec (annealing)
Step 4: 72°C, 10 sec (extension)
<return to step 2, 35X>
Step 5: 72°C, 1 min (polishing)
Step 6: 4°C, 2 mins
Step 7: END

PCR products were electrophoresed on 2% agarose gels and imaged with GelRed-Prestain (Biotium #41011) on a gel imaging station (Analytik-Jena UVP Chemstudio).

### RNAscope *in situ* hybridizations

Adult mice (P50-60) were anesthetized (3% isoflurane), and Fluorogold (2% FG, 300 μL; Fluorochrome) was injected into the left diaphragm through a lateral laparotomy. After 10 days, animals were deeply anesthetized (4% isoflurane, 96% O_2_), and perfused through the ascending aorta with 20 ml of 0.1 M phosphate-buffered saline (PBS; pH 7.4), followed by 20 ml of 4% paraformaldehyde (PBS, pH 7.4; Electron Microscopy Sciences, Fort Washington, PA). Spinal cords were dissected and stored in the perfusion fixative at 4°C for 4 hrs, then transferred to 20% sucrose for 8 hrs. Coronal sections (25 μm) were cut using a cryostat (Leica CM1860), mounted into glass slides, and stored at −80°.

We utilized the RNAscope™ Multiplex Fluorescent Reagent kit (Advanced Cell Diagnostics) ^10^, following the manufacture’s protocol. In brief, sections were incubated in PBS for 5 mins, followed by a 10 min incubation with H_2_O_2_ (ACD #322381) at room temperature (RT). This was followed by three washes with PBS (2 mins). Slides then treated with Protease IV (ACD #322381) for 30 mins (RT), followed by a 2 min PBS wash. Sections were then hybridized with RNAscope probes specific for *Kcnh1* (ACD #511141), *Kcnh2* (ACD #497681), *Kcnh5* (ACD #497691), *Kcnh6* (ACD #497701) and *Kcnh7* (ACD #1007281) for 2 hrs at 40°C, followed by a 2 min PBS wash. Following probe hybridizations, slides underwent sequential incubations with RNAscope Multiplex Detection Reagents (ACD #323110) consisting of FL v2 Amp 1, for 30 mins at 40°C, with a 5 min PBS wash following each step. Slides were then incubated with RNAscope Multiplex FL for 30 minutes at 40°C, followed by a 5 min PBS wash. To develop the HRP-C1 signal, sections were incubated with RNAscope Multiplex FL v2 HRP-C1 for 15 mins at 40°C, followed by a 5 min PBS wash. Finally, sections were treated with TSA Plus fluorescein (FP1168; amplification reagent, 1:250) for 30 mins at 40°C, followed by a 5 min PBS wash. Slices were then incubated with RNAscope Multiplex FL v2 HRP blocker for 15 mins at 40°C, followed by a 2 min PBS wash. Choline acetyltransferase (ChAT) was immunodetected using a primary polyclonal goat anti-ChAT antibody (#AB144P; Millipore; 1:100) diluted in PBS containing 2% normal donkey serum (#017-000-121, Jackson Immuno Research Laboratories) and 0.3% Triton X-100, incubated for 24 hrs at 4°C. Sections were then washed in PBS and incubated for 2 hrs with an Alexa 647-conjugated donkey anti-goat secondary antibody (#A-21447; Invitrogen; 1:500). All slides were mounted with Fluoromount (#00-4958-02; Thermo Fisher), and coverslips were sealed with nail polish.

Images were acquired on an Olympus VS120 slide scanner microscope, using Olympus VS-ASW-S6 software. Quantitation for cells positive for each RNAscope probe was achieved by manual counts of 55-94 phrenic motoneurons (ChAT^+^/FG^+^) from 8 selected sections, on slides representing each of 5 hybridization experiments, for an average of 588 phrenic motoneurons examined per hybridization.

### *In vitro* spinal cord slice preparation and recording

Coronal spinal cord slices containing the phrenic motor nucleus were prepared from neonatal mice between P4-12. Cholera toxin B (CTB) conjugated to Alexa Fluor-594 (2% in sterile PBS) was injected into the diaphragm of mouse pups at four sites using a 33-gauge needle attached to a Hamilton syringe. A volume of 0.5 µL was delivered per site over 1–2 minutes to ensure even labeling across the diaphragm. Partial brainstem and spinal cords were dissected following transcardio perfusion and rapid decapitation. Following extraction, the dorsal surface was glued onto an agar block cut at ∼15° angle. Neural preparations were sectioned stepwise at 300 μm in the transverse plane (Leica VT100S) and approximately 2 sections containing spinal cord segments C3-5 were saved for recording. Perfusion, dissection, and tissue sectioning occurred in ice cold artificial cerebrospinal fluid (aCSF) (in mM: 118 NaCl, 3.0 KCl, 25 NaHCO_3_, 1.0 MgCl_2_, 1.5 CaCl_2_, 30 D-glucose) equilibrated with carbogen (95% O_2_, 5% CO_2_), with an osmolarity between 305-312 mOsm. pH remained 7.4 ± 0.05 when equilibrated with carbogen.

Slices were placed in a recording chamber (Warner Instruments) with circulating aCSF (15 ml/min, 30°C) for study. The visual patch-clamp approach was used to record activity from single neurons. Recording electrodes were pulled from borosilicate glass (4-8 MΩ tip resistance) using a P-97 Flaming/Brown micropipette puller (Sutter Instrument Co., Novato, CA), filled with intracellular patch electrode solution containing: (in mM) 140 potassium gluconate, 1 CaCl_2_, 10 EGTA, 2 MgCl_2_, 4 Na_2_ATP, and 10 HEPES (pH 7.2, osmolarity: 309-314 mOsm). Neuronal activity was recorded under current-clamp and voltage-clamp protocols in whole-cell configuration, using video enhanced Dodt-IR optics and fluorescence for red fluorescent protein (RFP) on a Zeiss Axio Examiner.A1 microscope with a 40X water-immersion objective.

Electrophysiological recordings were made using an Axon 700B amplifier and pClamp v10 software (Molecular Devices, Sunnyvale, CA, USA), with signals digitized at 10 kHz and filtered at 2 kHz. Data analysis was performed using pClamp (Molecular Devices), LabChart (AD Instruments), and custom Python scripts within JupyterLab.

### *In vivo* electromyographic recordings

Adult mice were initially anesthetized using 3% isoflurane, followed by administration of urethane (1.5 mg/kg, intraperitoneally). During the experimental procedures, mice were placed in a supine position on a heated surgical table, custom-built to maintain their body temperature at 37°C. Mice were fitted with a nose cone, inhaling 100% oxygen throughout the experiment. Diaphragmatic muscle activity was monitored using electromyography (EMG) electrodes (A-M Systems) inserted into the lateral diaphragm. EMG signals were amplified by 10K X, filtered (300 Hz high-pass, 20 kHz low-pass), and analyzed for respiratory frequency and muscle activity throughout the duration of each respiratory cycle (AD Instruments. All pharmacological agents were delivered intraperitoneally. Morphine was administered at a dose of 150 mg/kg from a 10 mg/ml stock solution (Patterson Vet), and fentanyl at 500 µg/kg from a 50 µg/ml stock solution (Hikma). The doses for morphine and fentanyl were selected to achieve comparable levels of respiratory rate suppression. All Experimental trials were terminated by an overdose of isoflurane, followed by rapid decapitation.

## Acknowledgements

This study was supported by funding from the US National Institutes of Health grants R01 HL151389, R01 HL126523, R01 HL144801, P01 HL090554 (to J-M. R.), K99HL168211 (to N. J. B.), and the Brazilian Sao Paulo Research Foundation grant FAPESP 2021/05299-0 (to T. S. M.). We are grateful for comments by Drs. Phil Morgan, John P. Welsh, Fernando Pena, and Ryan Budde.

---

**Fig. S1. G/V plots for Kv10.1, Kv10.2, Kv11.1 and Kv11.3 measured under control, fentanyl (10 μM) and wash conditions.** Fentanyl exhibited a voltage-dependent and subtype-specific block of conductance, with a ∼70% block of *Erg*-subtypes (Kv11.1, Kv11.3). *Eag*-subtypes exhibited weaker fentanyl block (∼50%, Kv10.1; ∼20%, Kv10.2). All *Eag/Erg*-subtypes exhibited rapid block and variable partial recovery with wash. All plots shown as means with SEM, without fitting. See Methods for details of deriving conductance (G).

**Fig. S2. Molecular docking sites for fentanyl in Kv10.1(KCNH1), Kv10.2(KCNH5) and Kv11.1(KCNH2), predicted by AutoDock-Vina. a.** Side view of Kv10.1, with S463 at the base of the pore α-helix and Y491 at the top of S6 colored in blue. Fentanyl predicted to dock within the rings formed by these two residues contributed by each of the four subunits (left panel). A top-down view of this predicted Kv10.1-fentanyl docked structure through the K^+^ selectivity filter (colored in blue) shows the central piperidine ring of fentanyl, which is slightly electropositive, directly centered beneath the K^+^ selectively filter, in position to displace a focally coordinated K^+^ ion. **b.** Side view of Kv10.2, with T432 at the base of the pore α-helix and Y460 at the top of S6 colored in blue. Fentanyl predicted to dock below the ring formed by Y460, and in many conformations, consistent with its lower blocking ability for this *Eag*-subtype. **c.** Side view of Kv11.1, with S624 at the base of the pore α-helix and Y652 at the top of S6 colored in blue. Fentanyl predicted to dock within the rings formed by S624 and Y652, similar to Kv10.1, In addition, the phenylethyl ring of fentanyl is predicted to be aligned in close parallel orientation to the phenolic ring of Y652 (yellow circle), predictive of a strong stacked π-orbital interaction. These features of the predicted docking site may underlie the higher observed blocking ability of fentanyl for Kv11.1 and Kv11.3. Fentanyl is also predicted to dock asymmetrically relative to the central ion conduction pathway (right panel). See Methods for details of docking methods.

**Fig. S3. Amino acid sequence alignment of residues encoding the pore α-helix and S6 for all subtypes of the EAG superfamily of K^+^ channels subunits.** Highlighted in yellow are Kv11.1 residues (S624, Y652, F656) shown to mediate binding of the HERG specific blocker MK-499. *Eag-* and *Erg-* subtypes share greater conservation of these residues, compared to *Elk-*subtypes.

**Fig. S4. Morphine minimally alters spinal cord motoneuron excitability.** Bath application of 300 µM morphine under synaptic isolation induced negligible changes in the excitability of spinal motoneurons located in the ventral horn of the C3–5 spinal segments. Similarly, no significant changes in input resistance were noted following morphine application, suggesting that morphine has minimal direct effects on the intrinsic electrophysiological properties of these motoneurons. Two additional motoneurons showed inhibition of excitability with morphine (not shown).

**Fig. S5. *Eag/Erg* transcripts are widely expressed in mouse cortical areas from the Allen Brain Atlas. a.** Heatmap representation of single-cell transcriptomic data from the mouse cortex displayed for the two *Eag*-subtype genes (*Kcnh1, Kcnh5*) and the three *Erg*-subtype genes (*Kcnh1, Kcnh6, Kcnh7*), along with the μ-opioid receptor (*Oprm1*) and the four GIRK potassium channel subunits (*Kcnj3, Kcnj6, Kcnj9, Kcnj5*), and marker genes for inhibitor and excitatory neurons. Expression for *Eag/Erg* transcripts is strongly enriched for *Kcnh1*(Kv10.1) and *Kcnh7* (Kv11.3), whose expression is broadly seen in multiple transcriptomically-defined cell types, represented by both excitatory and inhibitory neuronal cell types. Particularly strong and consistent expression of *Kcnh7*(Kv11.3) is seen in *Pvalb* interneurons. Interestingly, *Oprm1* expression is relatively more selective and not as broadly seen throughout the cortex, suggesting that a large fraction of neuronal cell types expressing *Kcnh1*(Kv10,1) or *Kcnh7*(Kv11.3), may not also express molecular components necessary for inhibiting excitability by fentanyl through the known *Oprm1* Gα_i/o_ signaling cascade. **b.** *In situ* hybridization data for *Kcnh1*(Kv10.1) and *Kcnh7*(Kv11.3) seen in coronal slices from an adult mouse brain. Strong signals from both genes are observed in multiple layers of the cortex and in the hippocampus. *Kcnh1*(Kv10.1) is strongly expressed in the dentate gyrus, CA3 and CA2, whereas *Kcnh7*(Kv11.3) is strongly and preferentially expressed in CA1.

