## Supplementary figures and images for "Fentanyl blockade of K^+^ channels contribute to Wooden Chest Syndrome"

### Suppl-Wei et al-WCS

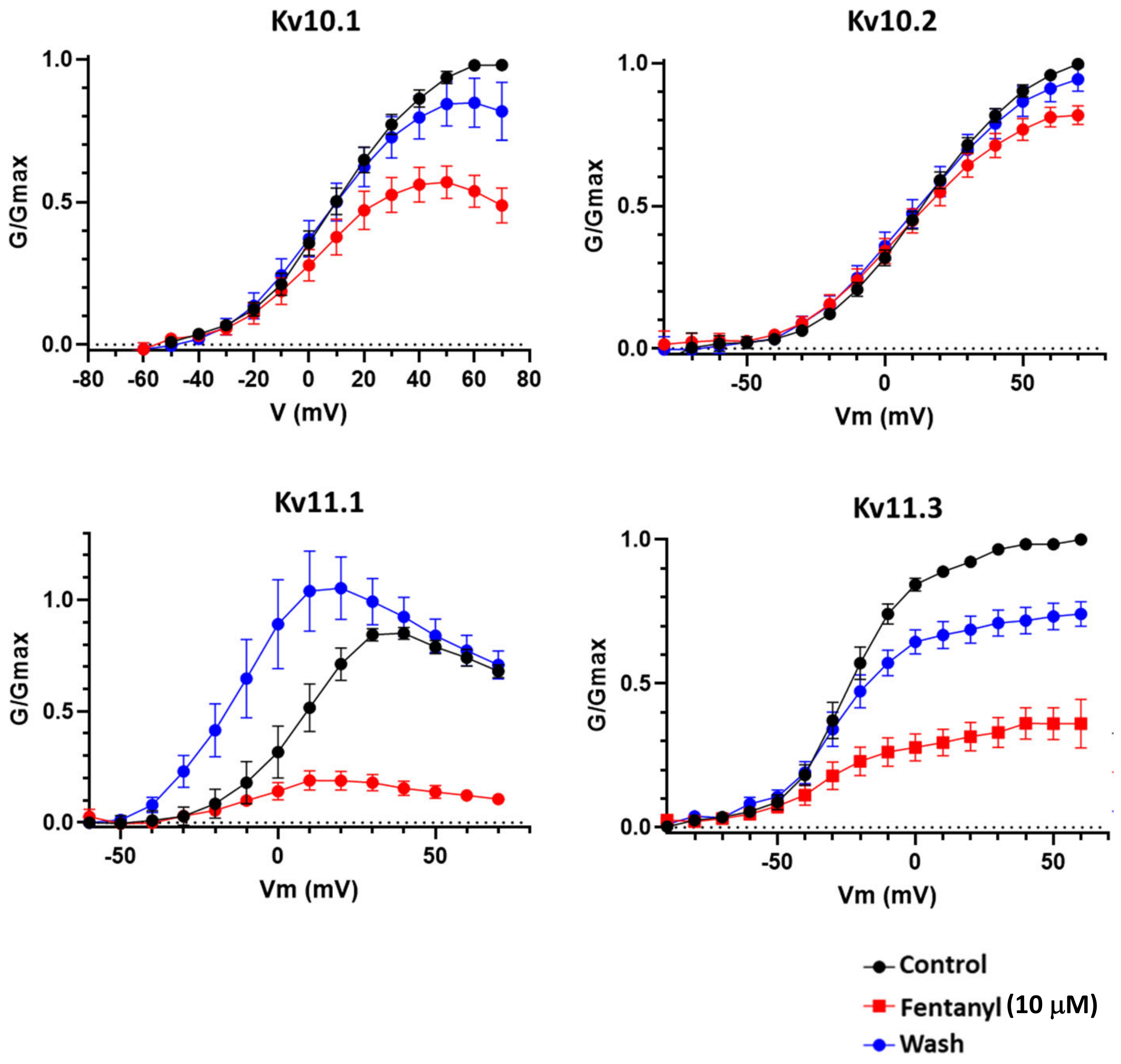

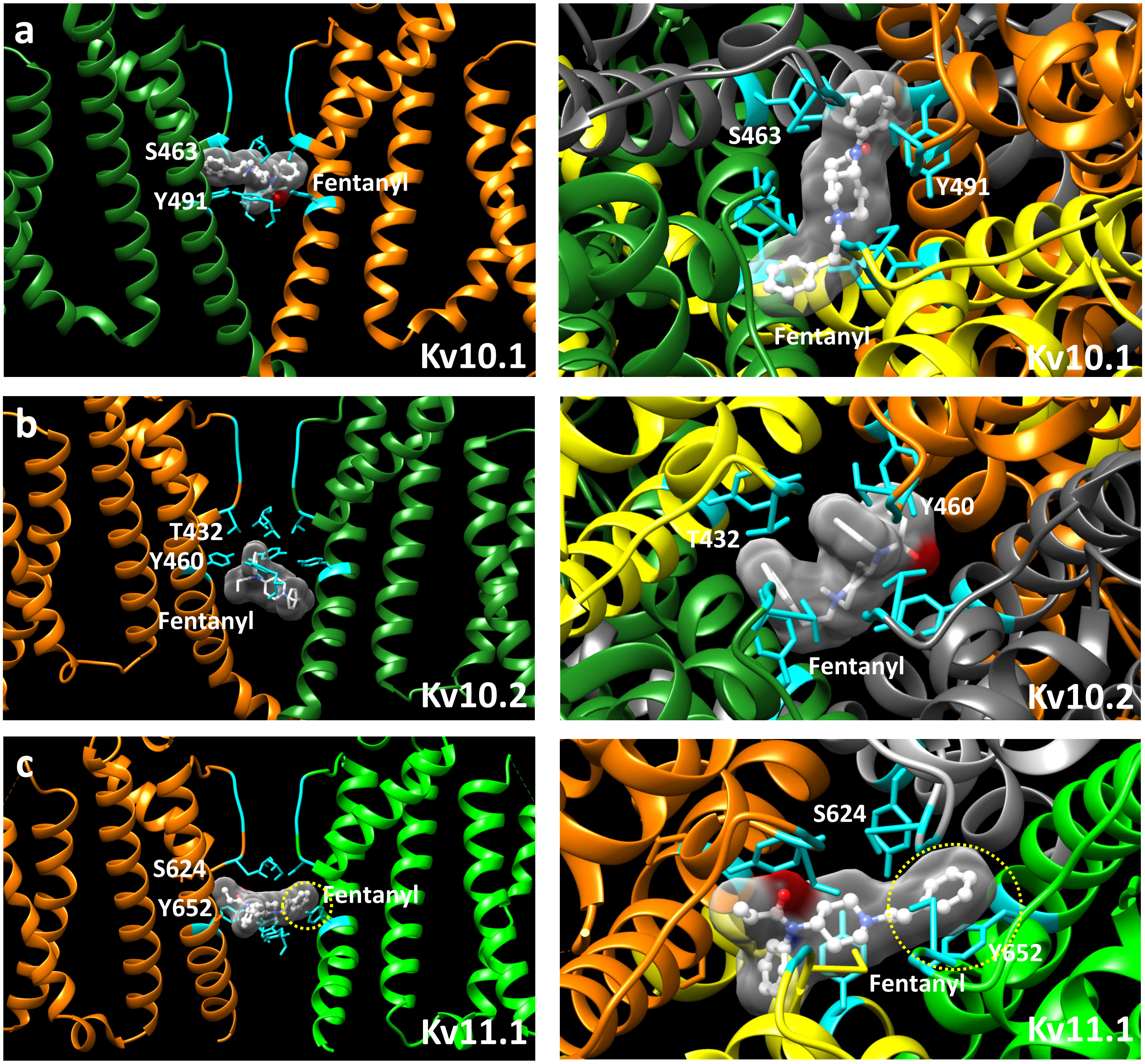

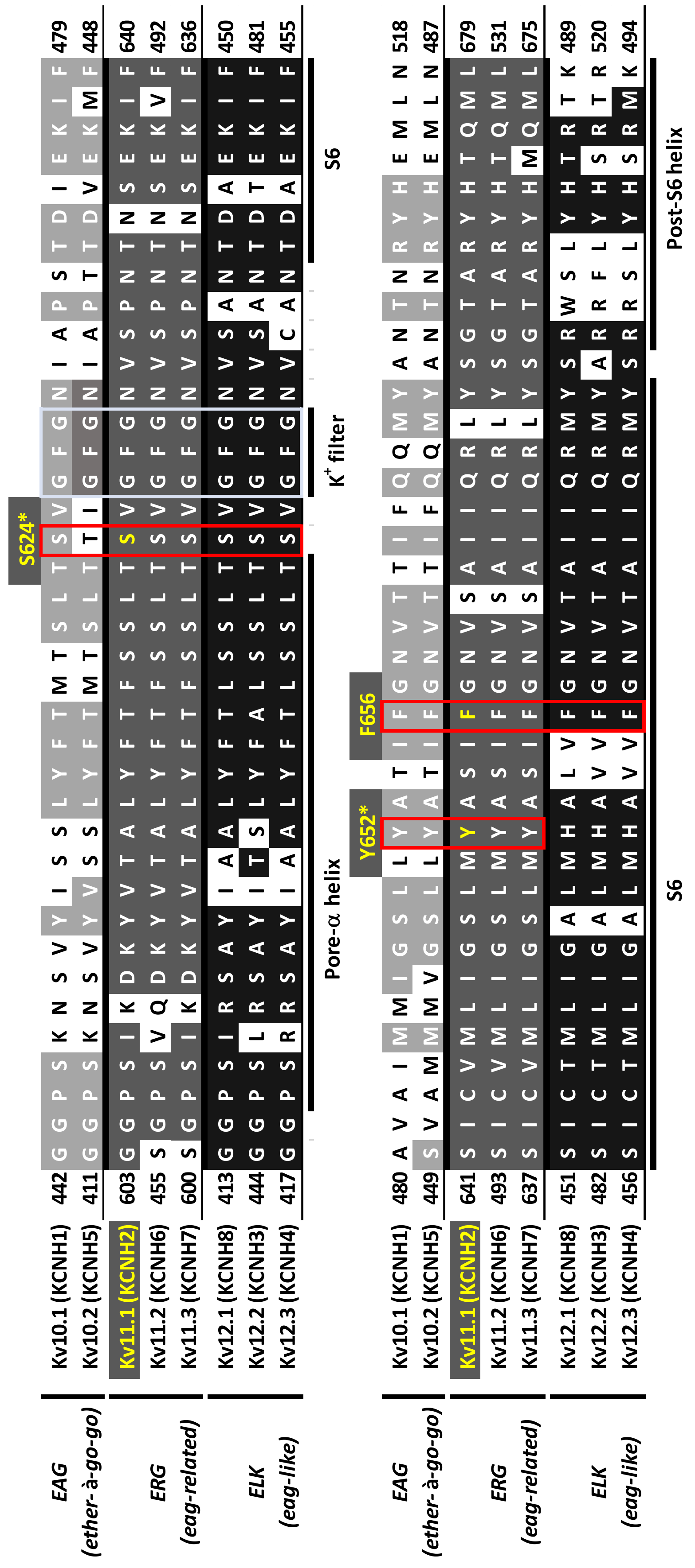

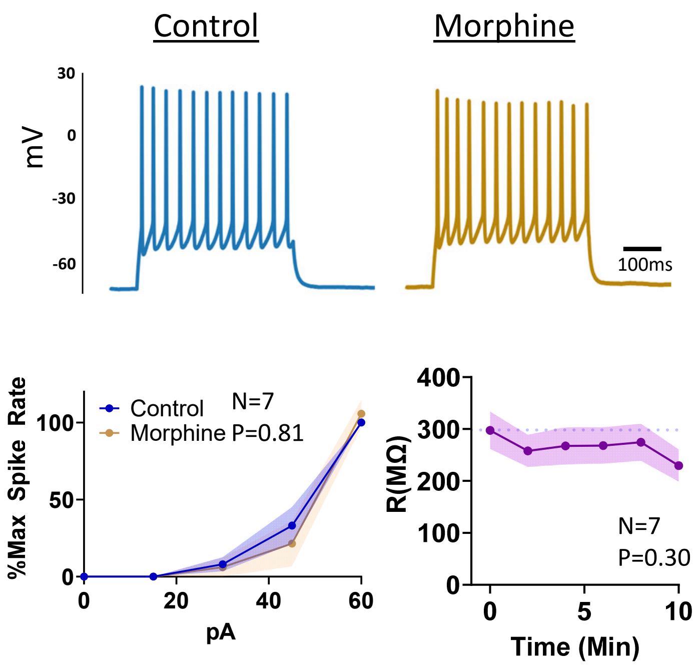

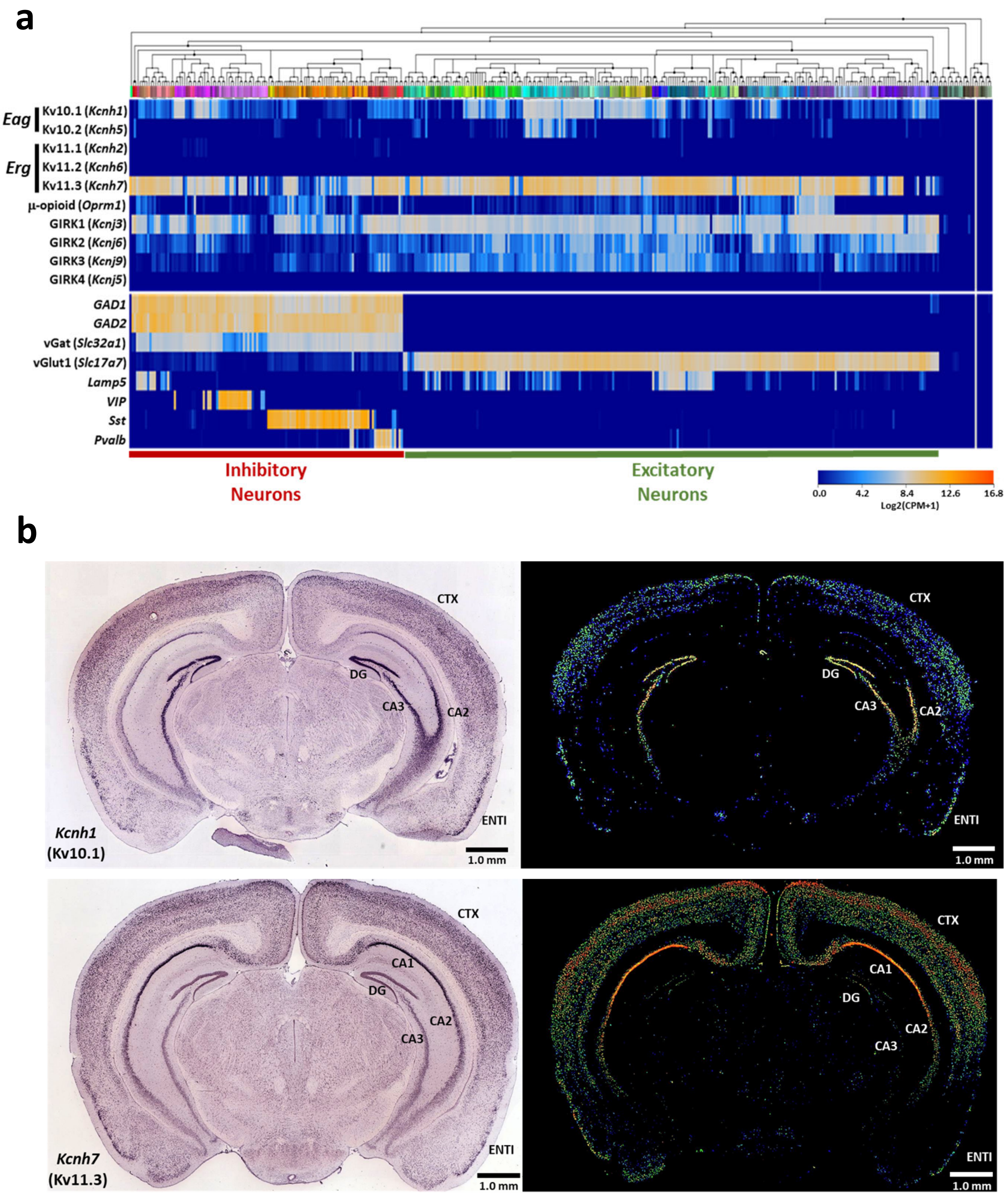


*(data from the Allen Mouse Brain Atlas)*

Fig. S5
